# RNA polymerase organizes into distinct spatial clusters independent of ribosomal RNA transcription in *E. coli*

**DOI:** 10.1101/320481

**Authors:** Xiaoli Weng, Christopher H. Bohrer, Kelsey Bettridge, Arvin Cesar Lagda, Cedric Cagliero, Ding Jun Jin, Jie Xiao

## Abstract

Recent studies have shown that RNA polymerase (RNAP) is spatially organized into distinct clusters in *E. coli* and *B. subtilis* cells. Spatially organized molecular components in prokaryotic systems imply compartmentalization without the use of membranes, which may offer new insights into pertinent functions and regulations. However, the function of RNAP clusters and whether its formation is driven by active ribosomal RNA (rRNA) transcription remain elusive. In this work, we investigated the spatial organization of RNAP in *E. coli* cells using quantitative superresolution imaging. We observed that RNAP formed large, distinct clusters under a rich medium growth condition and preferentially located in the center of the nucleoid. Two-color superresolution colocalization imaging showed that under the rich medium growth condition, nearly all RNAP clusters were active in synthesizing rRNA, suggesting that rRNA synthesis may be spatially separated from mRNA synthesis that most likely occurs at the nucleoid periphery. Surprisingly, a large fraction of RNAP clusters persisted under conditions in which rRNA synthesis was reduced or abolished, or when only one out of the seven rRNA operons (*rrn)* remained. Furthermore, when gyrase activity was inhibited, we observed a similar rRNA synthesis level, but multiple dispersed, smaller rRNA and RNAP clusters occupying not only the center but also the periphery of the nucleoid, comparable to an expanded nucleoid. These results suggested that RNAP was organized into active transcription centers for rRNA synthesis under the rich medium growth condition; their presence and spatial organization, however, were independent of rRNA synthesis activity under the conditions used but were instead influenced by the structure and characteristics of the underlying nucleoid. Our work opens the door for further investigations of the function and molecular nature of RNAP clusters and points to a potentially new mechanism of transcription regulation by the spatial organization of individual molecular components.

## Introduction

Prokaryotes are traditionally viewed as bags of freely diffusing enzymes. This view is rapidly changing. New studies now document that bacteria cells possess a remarkable degree of spatial organization of cellular components and activities without the use of membranes, offering a level of functionality and regulation previously underappreciated^1-4^. In both *E. coli* and *B. subtilis* cells grown in rich media, RNA polymerase (RNAP), the only enzyme responsible for all RNA transcription, was found to form dense foci instead of distributing homogenously within the cell^5,6^. Because the majority of cellular RNAP is dedicated to ribosomal RNA (rRNA) synthesis in fast-growing cells^7^, the transcription factory model was proposed^8^. This model suggests that dense RNAP foci are clusters of hundreds of RNAP molecules actively engaged in rRNA transcription, and that their formation is driven by active rRNA synthesis in fast-growing cells under optimal growth conditions (such as LB, 37°C)^5,8,9^. This prokaryotic transcription factory model is reminiscent of the RNAP I transcription factory model in eukaryotic cells, in which RNAP I forms concentrated, membrane-free condensates in the nucleolus for rRNA transcription^10,11^.

Understanding how and why RNAP is spatially organized in bacterial cells is important as this information could provide new insights into the mechanisms of transcription regulation in a complex, heterogeneous cellular environment. However, partially due to technical limitations in dissecting the subcellular organizations of small bacterial cells, many essential aspects of the bacterial transcription factory model remain elusive. In particular, despite a number of recent studies that extensively investigated the distribution and characteristics of RNAP clusters in *E. coii^12~u^*, whether RNAP clusters observed in fast-growing cells are indeed active in rRNA transcription, and whether RNAP clusters only form in the presence of active rRNA transcription, have not been directly examined. Previous studies have shown that treating cells with rifampicin, a global transcription inhibitor^15^, largely abolished the appearance of RNAP foci^9,14,16^. However, it remains unclear whether this change was due to diminished rRNA transcription activity, or the associated nucleoid structural changes under the condition of global transcription inhibition^17,18^.

In this study, we characterized the spatial distributions of RNAP and newly synthesized rRNAs under different transcription conditions in *E. coli* cells using quantitative superresolution imaging. We found that while RNAP clusters were associated with nascent rRNA synthesis under our rich medium growth condition, a high level of rRNA transcription activity and the presence of multiple *rrn* operons were not required for the presence of RNAP clusters. Instead, perturbing the supercoiling state of the chromosome via gyrase inhibition led to a redistribution of RNAP and rRNA clusters. Our work suggests that the characteristics and structure of the chromosomal may play a larger role than rRNA transcription activity in dictating the spatial organization of RNAP, and hence suggests a new mechanism of transcription regulation by spatial organization.

## Results

### RNAP formed distinct clusters in cells growing in rich defined medium

To investigate the spatial organization of RNAP in *E. coli*, we used a strain in which the chromosomal *rpoC* gene encoding for the β’ subunit of RNAP was replaced by a photoactivatable fluorescent gene fusion, *rpoC-PAmCherry*^13,14,19^. We verified that the resulting RpoC-PAmCherry fusion protein was expressed in full-length (Supplementary Fig. S1a), was incorporated efficiently into the RNAP core enzyme complex (Supplementary Fig. S1b) and supported wild-type (WT)-like cell growth as the sole cellular source of β’ subunit (Supplementary Fig. S1c). Therefore, the spatial distribution and dynamics of the RpoC-PAmCherry fusion protein should be representative of the native RNAP core or holoenzyme. In the text below, we refer to this fusion protein as RNAP-PAmCherry for simplicity.

Using RNAP-PAmCherry, we performed single-molecule localization-based superresolution imaging^20^ on exponentially growing live cells in EZ Rich Defined Media (EZRDM) at room temperature (25 °C, cell doubling time = 73 ± 1 min, hereafter termed as the rich medium growth condition, Supplementary Fig. S1c) with a measured two-dimensional (2D) spatial resolution of ^~^ 50 - 60 nm (Supplementary Fig. S2). We observed clustered distributions of RNAP-PAmCherry in individual cells (Fig. 1a). These clusters were distinct but less punctate compared to what has been reported previously in cells of faster growth rates^12-14^. The averaged cellular distribution of all RNAP localizations displayed a two-lobed pattern with a clear cleft in the middle (Fig. 1b), similar to that of the nucleoid imaged using three-dimensional (3D) structured illumination superresolution microscopy (SIM, Supplemental Fig. S3a). Using a truly monomeric mEos3.2-fused RNAP fusion protein, we verified that the clustered distribution was not due to the weak dimerization property of PAmCherry (Supplementary Fig. S4a, b). Furthermore, we developed a stringent algorithm to eliminate false clusters caused by repeated localizations of same molecules due to the blinking of fluorophores^21,22^, and still observed a clustered RNAP distribution (Supplementary Fig. S4c). Note that all the data used in this work was processed using the algorithm to elimimate repeated localizations.

**Figure 1.**
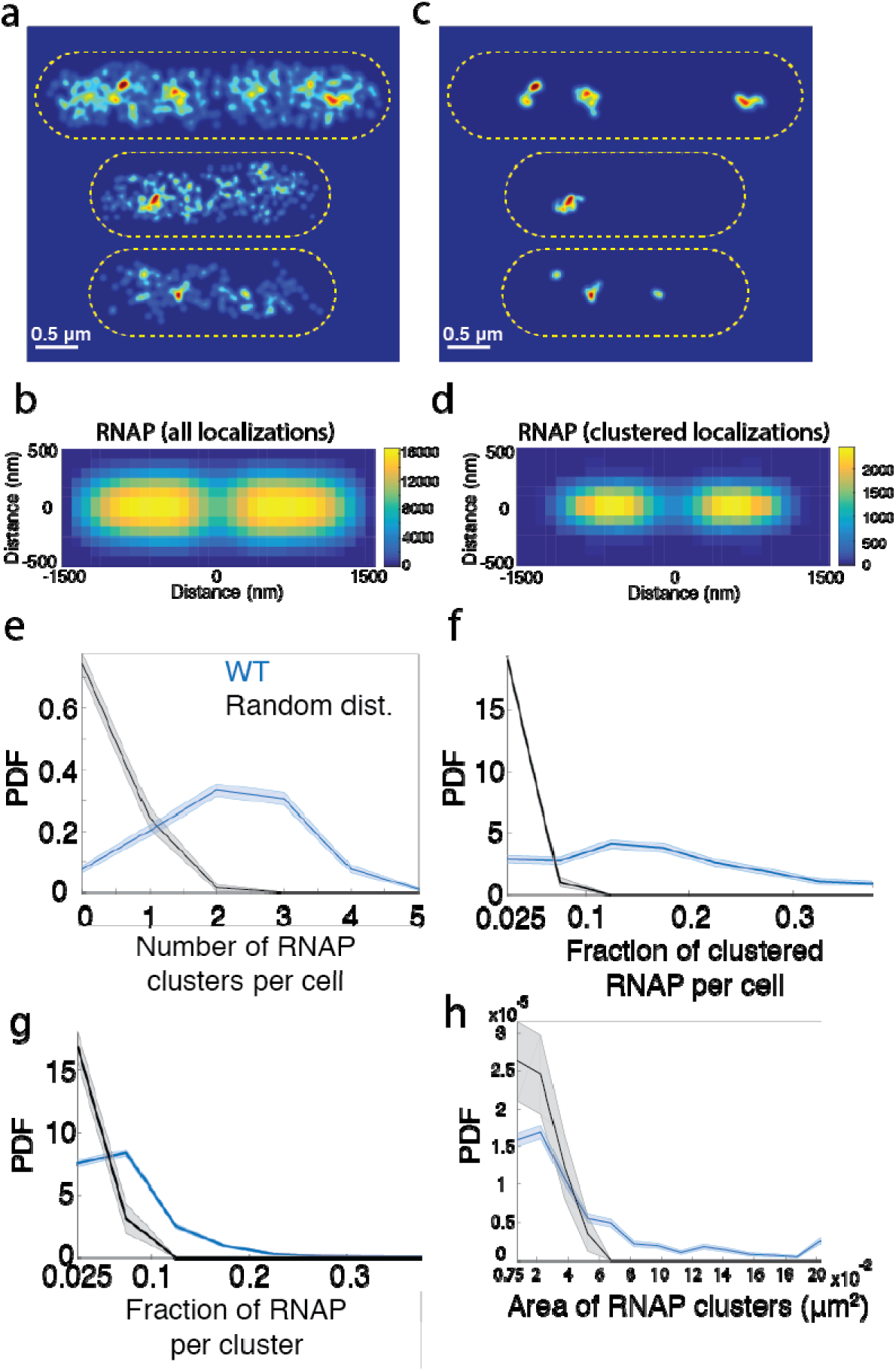
Quantitative characterization of RNAP clusters in live *E. coli* cells, (a) Representative superresolution images of RNAP (RpoC-PAmCherry) in three cells under the rich medium growth condition. Cell outlines are indicated in yellow dashed lines. Scale bar, 0.5 μm. (b) Two-dimensional (2D) histogram of all RNAP localizations in a standard 3 μm × 1 μm cell under the rich medium growth condition. Because of the symmetry of the cell shape in both long and short axes, we calculated the absolute displacement of each RNAP localization to the center of the cell, normalized its long axis displacement to the standard cell length, and duplicated the quartile cell histogram along both the long and short axes to produce a full-sized 2D histogram of RNAP distribution. The bin size of the 2D histogram is 100 × 100 nm. The color bar indicated localization numbers used in each bin. A total number of 564615 localizations of 664 cells are used to construct the 2D histogram, (c) Identification and isolation of RNAP clusters using a tree-clustering algorithm. RNAP clusters identified in the three cells in (a) are shown as examples, (d) 2D histograms of RNAP localizations in clusters as plotted in (b), a total number of 39438 localizations of 1385 RNAP clusters are used, (e) Distribution of the number of RNAP clusters per cell. The mean is 2.13 ± 0.05, μ ± s.e., n = 664. (f) Distribution of the fraction of clustered RNAP per cell. The mean is 0.16 ± 0.005, μ ± s.e., n = 664. (g) Distribution of fraction of RNAP localizations per cluster. The mean is 0.076 ±0.001, μ ± s.e., n = 1385. (h) Distribution of the area of RNAP clusters. The mean for the radius is 129 ± nm μ ± s.e., n = 1385 (assuming circularly shaped clusters). In all the graphs from (e to h), the blue curves are the experimentally measured distributions, and the black curves are those calculated from simulated random distributions using the same number of RNAP localizations in the same cell volume for all the cells. Shaded areas are standard errors calculated from bootstrapping. The average value of each graph is also summarized in Table S1.

To characterize RNAP clusters quantitatively, we performed a density-based threshold analysis to isolate individual RNAP clusters (Fig. 1c, Supplementary Table S1). The averaged cellular distribution of RNAP localizations inside these clusters also showed a similar, nucleoid-like pattern (Fig. 1d), but was more toward the center of the nucleoid compared to that of all RNAP localizations (Fig. 1b). On average, we detected ^~^2 dense RNAP clusters per cell (Fig. 1e). These clusters contained ^~^ 16% of total detected cellular RNAP-PAmCherry molecules (Fig. 1f), corresponding to approximately 350 RNAP molecules per cluster, given an average of ^~^ 5000 molecules of RNAP per cell (Fig. 1g, Supplementary Materials and Methods)^23,24^. On average these clusters occupied an area equivalent of that of a circle with a radius of ^~^ 130 nm (Fig. 1h). These properties were significantly different from what would be expected from a randomly distributed pattern of RNAP molecules (Fig. 1e-h, black curves, Supplementary Fig. S4a), confirming the clustering of RNAP in live *E. coli* cells under the rich medium growth condition.

### RNAP clusters colocalized with nascent rRNA synthesis sites in cells under the rich medium growth condition

To examine whether RNAP clusters are actively engaged in rRNA transcription, we probed the colocalization of RNAP clusters with nascent, or newly synthesized, rRNAs. We used a highly efficient fluorescence *in-situ* hybridization (FISH) probe labeled with Alexa Fluor 488 or 647 (Supplementary Fig. S5 and S6a) to target the 5’ leader region of the 16S precursor rRNA (pre-rRNA, Fig. 2a), which is absent from the mature 16s RNA inside the ribosome^25^. The 5’ leader degraded rapidly with a half-life of ^~^130 sec after being processed (Supplementary Fig. S6b); therefore, the FISH probe only identifies newly synthesized pre-rRNA. Using two-color superresolution imaging of pre-rRNA and RNAP-PAmCherry in fixed cells, we observed clear spot-like foci of pre-rRNA fluorescence signal with a spatial resolution of ^~^ 40 nm (Fig. 2b, Supplementary Fig. S2). On average we detected ^~^4 pre-rRNA clusters per cell containing more than 60% of total cellular rRNA localizations (Fig. 2c to f, Supplementary Table S2). Furthermore, we observed that qualitatively RNAP-PAmCherry clusters predominately coincided with these pre-rRNA clusters (Fig. 2b). To quantify the extent of spatial colocalization, we calculated the fraction of RNAP clusters that had any molecule from a pre-rRNA cluster within a radius ranging from 50 nm to 250 nm (half of the cell radius, Fig. 2g, blue curve, Material and Methods, Supplementary Fig. S7). We then compared the colocalization curve with the expected background level calculated by randomizing the positions of RNAP clusters in the same cells (Fig. 2g, black curve). We found that at all radii there were substantially higher fractions of RNAP clusters colocalizing with pre-rRNA clusters than that of the background level. For example, 83% ± 2% RNAP clusters (n = 404 RNAP clusters) had at least one pre-rRNA cluster within a radius of 50 nm (Supplementary Table S3). Given the significantly improved spatial resolution afforded by superresolution imaging, the high colocalization levels we observed between RNAP clusters and pre-rRNA clusters at a resolution limit (^~^ 40 – 60 nm) comparable to the moleuclar size of RNAP moleulces (^~^ 20 nm^26^) suggested that the majority of RNAP clusters were active in rRNA synthesis under the rich medium growth condition.

**Figure 2.**
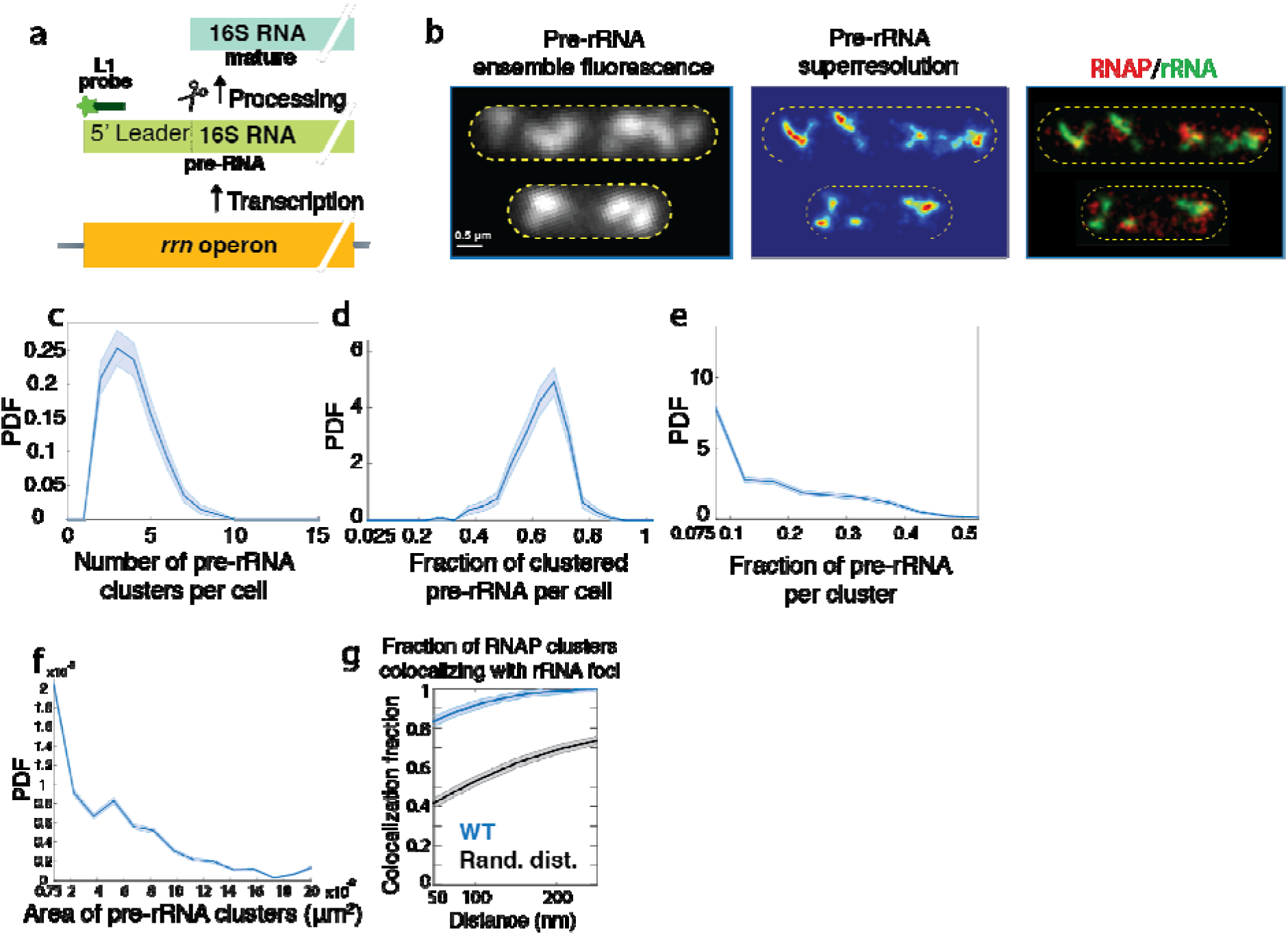
RNAP clusters colocalized with nascent pre-rRNA clusters under the rich medium growth condition. (a) Schematics of pre-rRNA detection. The dye-labeled L1 probe binds to the 5’ leader sequence of 16S rRNA that is cleaved off from mature 16S rRNA and rapidly degrades, (b) Left: ensemble pre-rRNA FISH images of cells (outlined in yellow) under the rich medium growth condition. Scale bar, 0.5 μm. Middle: representative pre-rRNA FISH superresolution images of two cells. Right: representative two-color superresolution images of RNAP-PAmCherry (red) and pre-rRNA FISH (green) of the two cells in the middle, (c) Distribution of the number of pre-rRNA clusters per cell. The mean is 3.86 ± 0.09, μ± s.e., n = 288. (d) Distribution of fraction of clustered pre-rRNA per cell. The mean is 0.63 ± 0.005, μ ± s.e., n = ± μ ± s.e., n = 288. (e) Distribution of fraction of pre-rRNA localizations per cluster. The mean is 0.16 ± 0.004, 1086. (f) Distribution of the area of pre-rRNA clusters. The mean for the radius is 127 ± nm μ ± s.e., n = 1086. In all the graphs from (c to f), the blue curves are the experimentally measured distributions. The average value of each graph is summarized in Table S2. (g) The fraction of RNAP clusters colocalizing with pre-rRNA clusters at different distances from 50 to 250 nm. The black curve is the simulated colocalization faction of RNAP clusters with pre-rRNA clusters when the spatial distribution of RNAP clusters was randomized in the same cells, and hence represented the basal level of colocalization due to chance. The plotted colocalization fraction is corrected for detection efficiency of pre-rRNA clusters (Supplementary Fig. S6a, S7), and all values are summarized in Table S3. In all the graphs the shaded areas are standard errors calculated from bootstrapping.

### RNAP clusters persisted in the absence of high levels of rRNA synthesis

To examine whether rRNA synthesis is the major driving force dictating the spatial organization of RNAP clusters as previously proposed^8,9,16,27^, we used drug inhibition to perturb rRNA transcription and subsequently observed the spatial organization of RNAP.

We first treated cells with rifampicin (RIF, 100 μg ml^−1^, 2 hours), a global transcription inhibitor that prevents transcription initiation but not elongation^15,28,29^. Consistent with previous studies^30^, we observed that rRNA synthesis was effectively abolished (Supplementary Fig. S8). The overall cellular distribution of RNAP exhibited a homogenous, single-lobed pattern without discernible central cleft, significantly different from that of untreated cells (Fig. 3a and b). However, RNAP clusters persisted, with a reduced number of RNAP clusters per cell and less RNAP molecules localized to clusters (^~^ 1.5 clusters/cell and 9%, respectively, Fig. 3c, d, Supplementary Table S1). These clusters also occupied a smaller area (^~^ 108 nm in radius) and contained ^~^ 20% fewer RNAP molecules compared to those of untreated cells (Fig. 3e, f, Supplementary Table S1).

**Figure 3.**
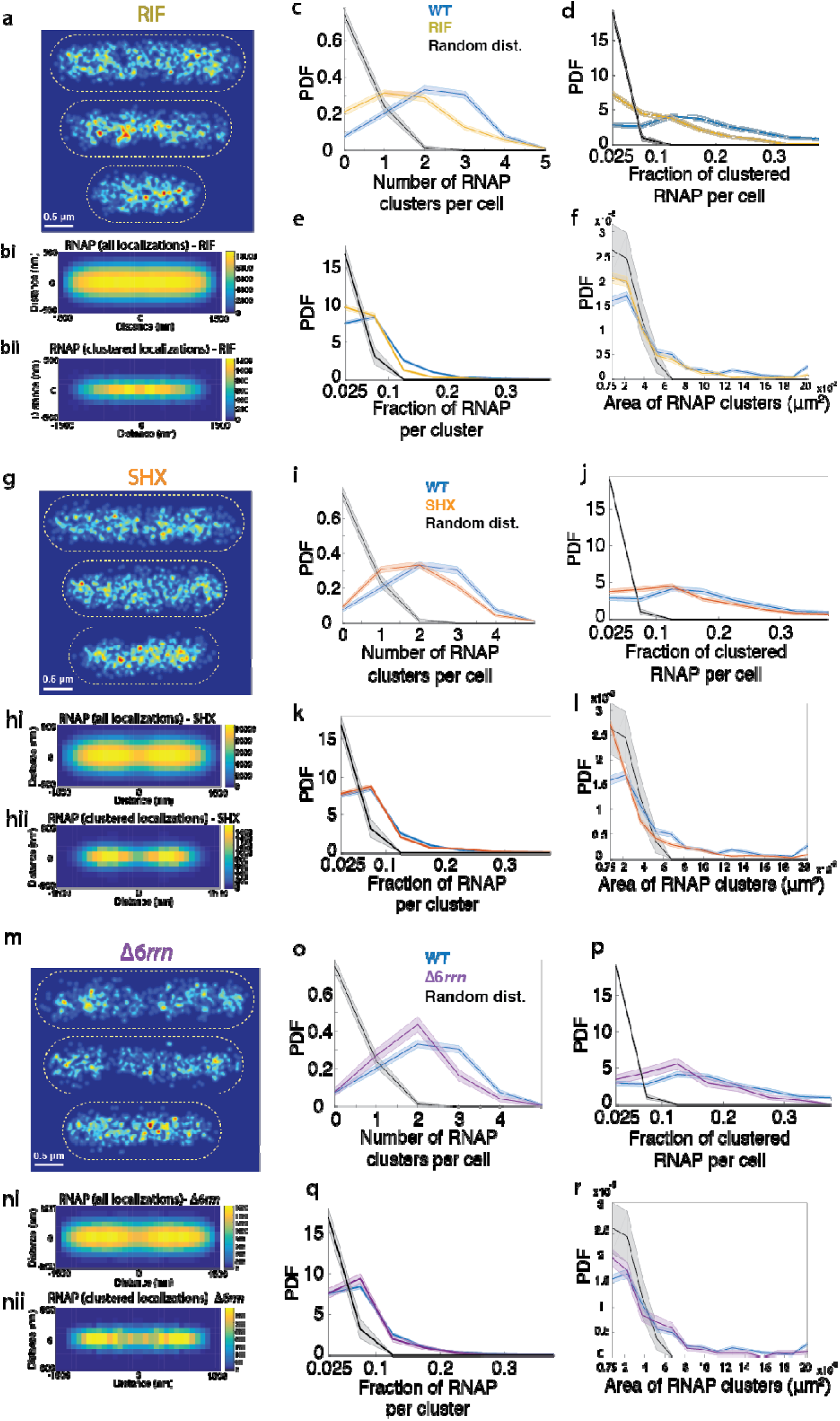
Characterization of RNAP clusters in live *E. coli* cells treated with rifampicin (RIF, top row, a to f), serine hydroxamate (SHX, middle row, g to l), and in a *rrn* deletion strain (Δ6*rrn*, bottom row, m to r). (a, g, m) Representative superresolution images of RNAP-PAmCherry. Scale bar, 0.5μm. (bi, hi, ni) 2D histogram of all RNAP localizations in a standard 3 μm × 1 μm cell, (bii, hii, nii) 2D histogram of only clustered RNAP localizations in a standard 3 μm × 1 μm cell. (c, i, o) Distribution of the number of RNAP clusters per cell. (d, j, p) Distribution of the fraction of clustered RNAP per cell. (e, k, q) Distribution of the fraction of RNAP localizations per cluster, (f, l, r) Distribution of the area of RNAP clusters. In (c-f, i-l, and o-r) the blue curves are those of the WT under the rich medium growth condition for comparison, and the black curves are those calculated from simulated random distributions using the same number of localizations in the same cell volume for all the cells under each condition. All the mean values of these graphs are summarized in Table S1. In all the graphs (c-f, i-l, and o-r), the shaded areas are standard errors calculated from bootstrapping.

We reasoned that because rifampicin inhibits all transcription and leads to global nucleoid expansion and likely rearrangement^17,18^, rifampicin may cause complicated secondary effects, leading to difficulties in the interpretation of cause and effect. Indeed, using SIM, we confirmed that cells treated with rifampicin showed a similarly expanded, homogenous distribution of the nucleoid compared to that of RNAP (Supplementary Fig. S3b, f). Therefore, we next treated cells with a relatively more specific rRNA syntheiss inhibitor, serine hydroxamate (SHX). SHX binds to seryl-tRNA synthetase, induces the stringent response and inhibits rRNA synthesis from the core *rrn* promoter p1^31-33^. As expected, we observed a dramatic reduction in total rRNA synthesis in SHX-treated cells as expected (^~^ 3% of untreated cells, Supplementary Fig. S8). When compared to rifampicin-treated cells, there was less noticeable expansion of the nucleoid as visualized using SIM (Supplementary Fig. S3c), and the total nucleoid volume of these cells showed no significant difference compared to WT cells (Supplementary Fig. S3f). However, RNAP clusters persisted and shared a similar, two-lobed distribution of the nucleoid (Fig. 3g, h, Supplementary Fig. S3c). The number of RNAP clusters per cell decreased (^~^1.9 clusters/cell, Fig. 3i,j, Supplementary Table S1), and their sizes were smaller (^~^ 104 nm, Fig. 3l), but they contained similar numbers of RNAP molecules compared to those in untreated cells (Fig. 3k). These results suggested that a high level of rRNA synthesis as that in the rich medium growth condition was not necessary for the spatial organization of RNAP clusters.

### RNAP clusters persisted in the presence of only one *rrn* operon per chromosome

Next, we reasoned that while rRNA transcription activity was diminished in cells treated with rifampicin or serine hydroxamate, it is possible that RNAP remained associated with multiple *rrn* operons that spatially colocalize with each other^34^, despite the lack of high transcription activity from these operons. To examine this possibility, we used a Δ*6rrn* strain, in which six out of seven *rrn* operons (except for *rrnC)* were removed from the chromosome^16^. The *Δ6rrn* strain grew at a slower rate than WT cells under the same rich medium growth condition (cell doubling time = 91 ± 1 min, Supplementary Fig. S9), and showed a significant reduction in total rRNA synthesis (^~^28% of WT cells, Supplementary Fig. S8). However, the spatial distribution of RNAP and the properties of RNAP clusters in the *Δ6rrn* strain were remarkably similar to those of SHX-treated cells (Fig. 3m to r, Supplementary Table S1), and remained highly colocalized to residual pre-rRNA clusters (Supplementary Fig. S10, Table S3). Additionally, we found that the nucleoid morphology and the total nucleoid volume of these cells were comparable to WT cells (Supplementary Fig. S3d, f). These results suggested that RNAP clusters did not require a high level of rRNA synthesis activity or the presence of multiple *rrn* operons.

### Inhibition of gyrase activity led to a redistribution of RNAP clusters and rRNA synthesis sites

Our results so far showed that under the rich medium growth condition, nearly all RNAP clusters are active in rRNA synthesis; however, neither a high level of rRNA synthesis activity nor the presence of multiple *rrn* operons was required for the formation of RNAP clusters. The question is then what would be responsible for spatially organizing RNAP into clusters. We noticed that in all conditiones we tested, the cellular distribution of RNAP closely mimicked that of the corresponding nucleoid structure visualized using 3D SIM imaging (Supplementary Fig. S3). These observations suggested that the spatial organization of RNAP might reflect that of the underlying nucleoid organization rather than the transcription activity or presence of multiple *rrn* operons.

The *E. coli* chromosome is highly compact and organized at different levels from topological domains to macrodomains (MDs)^35-37^. These organizations likely dictate the spatial arrangement of different DNA segments, upon which RNAP may preferentially bind and form clusters. Negative supercoiling is a major chromosome compacting factor, and it is only introduced by gyrase, a type II topoisomerase in *E. coli^38^.* We thereby examined specifically the effect of DNA supercoiling on the spatial organization of RNAP.

We treated WT cells with a gyrase inhibitor novobiocin (NOV, 300 μg ml^−1^ for 30 min) and performed two-color superresolution imaging using pre-rRNA FISH and RNAP-PAmCherry. Novobiocin inhibits gyrase activity by abolishing ATP binding to the ATPase domain in the GyrB subunit^39,40^. We found that the average rRNA synthesis per cell was not significantly affected by the inhibitor, as the total intensity of pre-rRNA FISH signal remained similar to untreated cells (Supplementary Fig. S11a), even when high inhibitor concentrations and long-time treatment were used (Supplementary Fig. S11b). We further verified that the persistent rRNA synthesis during gyrase inhibition was not due to altered rRNA degradation kinetics in the presence of the inhibitor (Supplementary Fig. S11c). The minimal effect of gyrase inhibition on rRNA synthesis has been observed previously^41,42^, although conflicting results have been reported as well^43,44^. Interestingly, while the total pre-rRNA FISH signals remained unchanged under our experimental condition, we observed a greater number of smaller (on average ^~^ 6 per cell) and less dense (^~^9% of localizations) pre-rRNA clusters occupying a larger area of the cell compared to untreated cells (Fig. 4a to c and e to h). RNAP clusters persisted in these gyrase-inhibited cells (Fig. 4d and i to l), remained highly colocalized with pre-rRNA clusters (Fig. 4m), but contained fewer RNAP molecules. Interestingly, the cellular positioning of RNAP clusters and pre-rRNA clusters expanded ^~^ 50 nm toward the nucleoid periphery (Supplementary Fig. S12). As such, the cellular distributions of pre-rRNA and RNAP clusters exhibited a similarly, spatially dispersed pattern compared to untreated cells (Fig. 4c, d) and mimicked that of the expanded nucleoid under the same condition (Supplementary Fig. S3e, f). A different gyrase inhibitor, Nalidixic acid (NA, 50 μg ml^−1^ for 10 min), which acts on the GyrA subunit by stabilizing the DNA-cleaved complex, produced a similar effect (Supplementary Fig. S13). These results suggested that the characteristics and organization of the nucleoid, here in particular compacted by negative supercoiling, could play a larger role in the spatial distribution of RNAP clusters compared to rRNA transcription activity.

**Figure 4.**
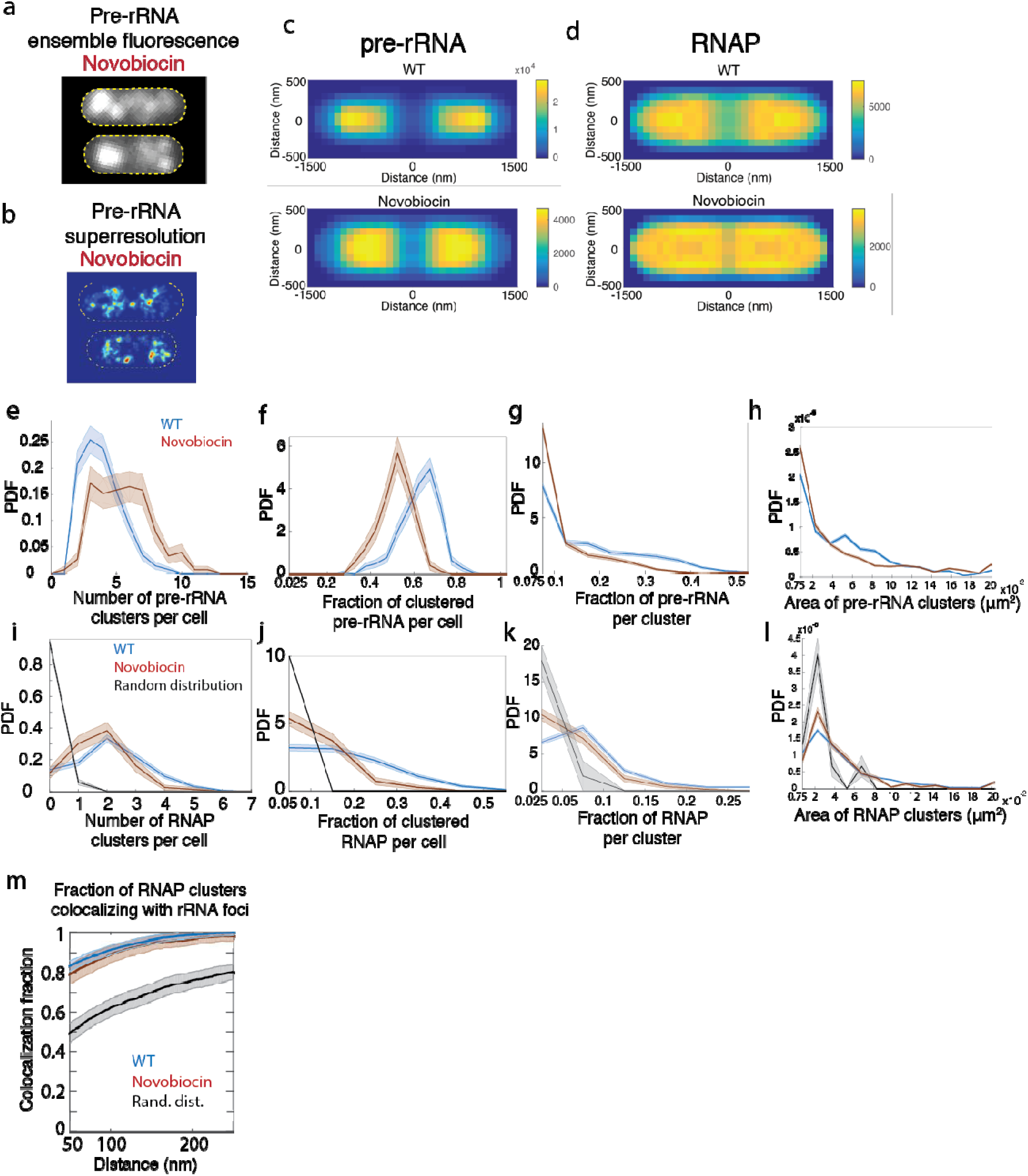
Inhibition of gyrase activity led to dispersed distributions of RNAP and pre-rRNA. (a) Ensemble fluorescence of Pre-rRNA FISH signal in fixed, novobiocin-treated cells. Individual cells are outlined in yellow. (b) Representative superresolution images of pre-rRNA distribution in fixed, novobiocin treated cells. (c) 2D histograms of all pre-rRNA localizations in a standard 3 μm − 1 μm fixed cell under the rich medium growth condition (top) and in cells treated with novobiocin (bottom). (d) 2D histograms of all RNAP localizations in a standard 3 μm × 1 μm fixed cell under the rich medium growth condition (top) and in cells treated with novobiocin (bottom). (e-l) Distributions of properties of pre-rRNA (e-h) and RNAP clusters (i-l) in Novobiocin-treated cells. (e, i): Distribution of the number of clusters per cell. (f, j): Distribution of fraction of clustered pre-rRNA (f) or RNAP (j) per cell. (g, k): Distribution of fraction of pre-rRNA (g) or RNAP (k) localizations per clusters. (h, l): areas of clusters. (m) Fraction of RNAP clusters colocalizing with pre-rRNA clusters in novobiocin-treated cells. In all plots the WT rich medium growth conditions are plotted in blue for comparison; novobiocin-treated conditions are in dark red, and the background colocalization levels using simulated images are in black. All shaded areas are standard error calculated using bootstrapping. All the mean values of these graphs are summarized in Table S2 and S3.

## Discussion

In this study, we investigated the transcription factory model in detail using a combination of quantitative superresolution imaging and perturbation analyses. We provided direct evidence demonstrating that under the rich medium growth condition the majority of RNAP clusters were actively engaged in rRNA transcription. These clusters’ presence and spatial distribution did not appear to require a high level of rRNA transcription activity or the presence of multiple *rrn* operons. Our results suggest that these clusters were instead likely influenced by the underlying nucleoid structure. Below we compare our results with previous work and discuss the implications of this work.

### Spatial organization of RNAP

Using quantitative superresolution imaging, we found that in *E. coli* cells grown in rich medium, RNAP was spatially organized into large, dense clusters occupying the same area as the nucleoid. These clusters had a radius of ^~^ 130 nm (Fig. 1g), and likely represented collections of multiple small RNAP clusters observed in a previous study^14^. Given a total cellular level of RNAP at ^~^5000 molecules per cell^23,24,45^ under a similar growth condition, and that majority (^~^90%) of cellular RNAP remain bound on DNA^46^, we estimated that each RNAP cluster contained ^~^ 350 molecules. The cellular distribution of all RNAP molecules exhibited a two-lobed pattern with a clear cleft in the middle (Fig. 1b), mimicking that of two replicated and segregated nucleoids (Supplementary Fig. S3). Compared to the distribution of all RNAP molecules, RNAP clusters were tighter and more concentrated toward the center of the two lobes with an average distance of ^~^ 75 nm from the center of the cell (Supplementary Fig. S14). This observation was consistent with previous superresolution studies of spatially separated ribosome and RNAP in *E. coli*^12^ — ribosome at the nucleoid boundary while RNAP predominately at the center, but different from the observation in another study that a significant portion of immobilized RNAP molecules localized to the periphery of the nucleoid^14^. The spatial segregation between ribosome and RNAP has been used to question the coupling between transcription and translation in bacterial cells^12,46^. It is possible that periphery-localized small RNAP clusters, which may be undetectable in our stringent distance-based clustering algorithm, could be RNAP molecules actively engaged in mRNA transcription that is coupled to translation by ribosome, while the nucleoid center-localized large RNAP clusters we observed were engaged in rRNA synthesis, which does not require translation. As such, these results suggested that rRNA transcription could be spatially separated from mRNA transcription. Additionally, it has been well documented that RNAP clusters are dynamic and sensitive to growth conditions^5^. Under our rich medium growth condition, cells grew at a slower rate compared to previous studies, which also likely contributed to the observed differences in the cellular distribution of RNAP clusters.

### Spatial organization of pre-rRNA clusters

We used a pre-RNA FISH probe targeting the leader sequence of the 16S rRNA to detect rRNA transcription activity. Because newly synthesized pre-rRNAs are processed before they are incorporated into ribosomes, the pre-rRNA probe marks new rRNA synthesis sites. Compared to RNAP, pre-rRNAs formed similarly sized (^~^130 nm in radius) but denser (containing > 60% of detected cellular pre-rRNAs) clusters. The overall cellular distribution of pre-rRNAs also exhibited a two-lobed pattern, but relatively more concentrated at the center of the nucleoid compared to RNAP clusters in fixed cells (Fig. 4c, d, Supplementary Fig. S12). On average we observed ^~^ four pre-rRNA clusters per cell or two per nucleoid. Because four out of the seven *rrn* operons reside close to the replication origin *oriC* on the chromosome, it is possible that the two pre-rRNA clusters reflected two copies of replicated *oriC* for each chromosome, and thus most cells contained two copies of the chromosome with four copies of *oriC* region, consistent with previous observations when the *Ori* region was labeled^47^. Alternatively, it is also possible that the two pre-rRNA clusters reflected two groups of transcribing *rrn* operons on the same copy of chromosome that are spatially distinguishble from each other under our resolutions. A recent study found that, except for *rrnC*, all the *rrn* operons are within a spatial distance of ^~^80 – 130 nm (median of distributions) to each other^34^, but it remains unknown whether these *rrn* operons indeed co-occupy the same area in the nucleoid and whether they could be accommodated in one pre-rRNA cluster (radius of ^~^ 130 nm). Because of the nearly identical pre-rRNA sequences of all the *rrn* operons, we could not design a probe with high confidence to distinguish the transcription activity of individual *rrn* operons, and hence further investigation is required to address whether each pre-rRNA cluster reflects the transcription activity from individual or a collection of *rrn* operons.

### Contribution of rRNA transcription activity to the spatial organization of RNAP

We observed that under the rich medium growth condition, RNAP clusters highly colocalized with pre-rRNA FISH probe signals (> 80% RNAP clusters colocalized with pre-rRNA clusters within 50 nm). Because the pre-RNA FISH probe only detects newly transcribed pre-rRNAs instead of mature, stable rRNAs (Supplementary Fig. S6b), and the two-color superresolution imaging offers spatial resolution (Supplementary Fig. S2) closer to the molecular size of RNAP complexes (^~^ 20 nm^48^) compared to conventional fluorescence imaging, we concluded that under the rich medium growth condition the majority of RNAP clusters were transcribing rRNAs actively. This result is the most direct demonstration of the activity of RNAP clusters. Previous studies have assumed but not validated that RNAP clusters are transcribing rRNAs in fast-growing cells^8,13^; in one study it was shown that RNAP clusters colocalized with clusters of two transcription factors NusA and NusB that are involved in anti-termination of rRNA synthesis and ribosome biogenesis^49^. In fast-growing cells, rRNA synthesis rate is at its maximum, and there is little pause or stalled RNAP in rRNA operons^50^. Under different growth conditions, however, Nus factors have also been implicated in associating with stalled/paused RNAP molecules^51,52^.

The ability to observe nascent pre-rRNA synthesis sites allowed us to examine whether rRNA transcription activity is the driving force for RNAP cluster organization under the conditions used. We blocked rRNA transcription to different degrees using rifampicin, serine hydroxamate, and a mutant strain lacking six of the seven *rrn* operons (Δ6*rrn).* We found that rifampicin-treated cells showed the most significant changes compared to others. The cellular distribution of all RNAP molecules in rifampicin-treated cells exhibited single, elongated lobes without discernible middle cell cleft. While this changed distribution could be due to the redistribution of RNAP on the nucleoid after the inhibition of rRNA synthesis, it also could be caused by the reorganization/expansion of the nucleoid, as the nucleoid exhibited similar changes in these cells (Supplementary Fig. S3b). Most importantly, cells still contained a significant number of RNAP clusters (1.5 per cell, Supplementary Table S1). Serine hydroxamate-treated cells were remarkably similar to Δ6*rrn* cells in all aspects of RNAP clusters: compared to WT cells, these cells exhibited two-lobed distributions of RNAP with less prominent but clear mid-cell cleft; the average number of RNAP clusters dropped minimally to ^~^ 1.9 per cell, and the remaining RNAP clusters had similar size and number of molecules (Fig.3g to 3r, Supplementary Table S1). However, rRNA transcription activity of Δ*6rrn* cells was less than one-third of that in WT cells, and SHX-treated cells had negligible rRNA synthesis (Supplementary Fig. 8). The drastically different rRNA transcription activities but similar organizations of RNAP under the two different conditions hence suggested that rRNA transcription activity may not be the driving force for the organization of RNAP clusters under our experimental conditions. Furthermore, because there was only one copy of the *rrnC* operon in the *Δ6rrn* strain, it is unlikely that multiple *rrn* operons were required for the formation of RNAP clusters as previously proposed^5^. Taken together, these results argued strongly against the hypothesis that rRNA transcription activity from multiple *rrn* operons contributes significantly to the spatial organization of RNAP. Consistent with this notion, a recent study reported that multiple *rrn* operons could colocalize with each other independent of rRNA transcription activity^34^.

### Contribution of nucleoid structure to the spatial organization of RNAP

In all of our experiments we observed that the cellular distribution of RNAP mimicked the shape of the underlying nucleoid irrespective of rRNA synthesis activity (Supplementary Fig. S3). We thereby turned to investigate the contribution of the nucleoid structure on the spatial distribution of RNAP by inhibiting gyrase. Gyrase is the only type II topoisomerase in *E. coli* that introduces negative supercoiling into the chromosome, which is the major force in compacting the nucleoid^38,53^. In gyrase-inhibited cells, we observed similar levels of pre-rRNA signal (Supplementary Fig. S11) but saw a significant shift in the cellular positioning of the RNAP clusters and pre-rRNA clusters, both of which expanded ^~^ 50 nm toward the nucleoid periphery (Fig. 4, Supplementary Fig. S12, S13). Previous studies have documented that gyrase inhibition affects the expression of more than three hundred mRNA genes that are sensitive to supercoiling^54,55^, but produced mixed results on the effect of rRNA transcription^42-44^. The pre-rRNA FISH probe detects the 5’ leader sequence of 16S rRNA, hence the unchanged FISH signal in gyrase-inhibited cells only reflected unaltered rRNA transcription initiation. It is possible that rRNA elongation was inhibited due to accumulated positive supercoiling ahead of transcription in the absence of gyrase. In such a case, we should expect that after a long inhibition time, the rRNA transcription initiation rate would gradually decrease as accumulated positive supercoiling eventually inhibits transcription initiation, which was demonstrated previously for the production of mRNA *in vitro^56^.* Nevertheless, we observed similar pre-rRNA signal even after we incubated cells with high concentrations of novobiocin (300 to 1200 μg ml^−1^) for extended periods of time (90 – 150 min, Supplementary Fig. S11), suggesting that total rRNA transcription activity in these cells was minimally affected. Therefore, these experiments most likely suggested that compared to rRNA transcription activity, the nucleoid structure contributes more significantly to the spatial organization of RNAP. It is possible that a relaxed chromosome repositioned different DNA segments (upon which RNAP clusters form) to occupy a larger cell volume, as suggested by SIM imaging of the gyrase-inhibited nucleoid (Supplementary Fig. S3e, f). Clearly, further studies, such as genetic and biochemical perturbations of nucleoid-organization factors, are required to investigate the effect of the nucleoid structure on the spatial organization of RNAP.

In summary, our study demonstrated that there was a rRNA transcription activity-independent spatial organization of RNAP in *E. coli* and that the underlying nucleoid structure likely played an important role in organizing RNAP clusters. Evidently, further experiments investigating the molecular nature and function of RNAP clusters are required. In particular, we do not know whether these RNAP clusters we observed are associated with specific chromosomal DNA sequences, or whether they are self-promoting oligomeric complexes similar to liquid droplets observed in eukaryotic cells, which are mediated by multivalent interactions among proteins and nucleic acids^57,58^. Intriguingly, it has been shown that a small regulatory ncRNA 6S can sequester σ70 holoenzyme of RNAP and help *E. coli*’s rapid transition into stringent response conditions; these RNAP-RNA complexes may also contribute to clusters we observe under conditions where transcription activity from σ70 promoters is low^59,60^. Furthermore, we do not know the biological significance of RNAP clusters. In eukaryotic cells, it was suggested that RNAP clusters might represent pre-formed transcription complexes that are “poised” ready for rapid transcription induction^61-64^. In bacterial cells, such a role has not been demonstrated, but studies have shown that there are typically higher levels of RNAP association at promoter and promoter-like sequences than within coding sequences^65-70^. Perhaps looking at the colocalization of RNAP with important transcription regulators (NusA^71,72^, NusB^73,74^, NusE^75^, NusG^65,76^, and SuhB^77^, *etc.*) that interact with RNAP under different conditions would help elucidate the molecular makeup and functional significance of RNAP clusters. Regardless, further investigations into the spatial organization of RNAP in small bacterial cells will certainly bring in new knowledge complementing *in-vitro* biochemical and *in-vivo* genetic studies of prokaryotic transcription.

## Materials and Methods

### Bacterial strains and constructions

All bacterial strains used in this study were listed in Supplementary Table S4. The wild-type (WT) strain background was MG1655. The RpoC-PAmcherry (CC253) and RpoC-mEos3.2 (XW023) chromosomal fusion strains were constructed using λ-RED-mediated homologous recombination^78^. Specifically, the linear fragment containing the fluorescent protein ‘*FP-frt-kan^R^-frt’* sequence was first generated and subcloned into the pKD13 plasmid^78^. The 50-bp homologous flanking regions were then added to the linear fragment using primer pairs 15-16 and 17-18. The linear fragment was transformed into MG1655 cells containing the pKD46 plasmid with 0.2% L-arabinose induction. Recombinants grown on LB + kanamycin plates were verified by colony PCR. The pKD46 plasmid was next cured by growing cells at the restrictive temperature 37 °C. The RpoC-PAmCherry fusion in the Δ6*rrn* strain (CC302) was constructed similar to described above^16^. In later constructions, the *frt-kan^R^-frt* cassette was flipped out using the PCP20 plasmid^78^ to generate strain ACL002. A chromosomal DNA site marker (tetO^6^) was inserted at different chromosomal locations of ACL002 to generate a series of dual-labeled strains (ACL066, ACL020, XW030, XW033, ACL036, XW017, and XW018) using primer pairs 1 to 14, and λ-RED mediated homologous recombination as described above. A plasmid expressing the TetR-EYFP reporter was constructed from pZH102R33Y29^79^ and transformed into all the dual-labeled strains. We imaged both RNAP and DNA localizations of all the strains in live cells, but only included RNAP localization data in this work due to the limit of space. DNA localization data will be described in an accompanying study.

### Cell growth

Single *E. coli* colonies were picked from freshly streaked LB plates and cultured in EZ Rich Defined Media (EZRDM, Teknova) with 0.4% glucose, at room temperature (RT) overnight with shaking. Antibiotics (kanamycin (Sigma-Aldrich 1355006) and carbenicillin (Sigma-Aldrich C3416)) were added at 50 μg ml^−1^ when appropriate. The next morning, cells were reinoculated into fresh EZRDM with 0.4% glucose and grown at RT until they reached mid-log phase (O.D._600_ ^~^ 0.3-0.4). To induce TetR-EYFP expression, cells were harvested and resuspended in fresh EZRDM supplemented with 0.3% L-arabinose and 0.4% glycerol and allowed to grow for two additional hours (this condition was used for all live cell imaging experiments reported in this work). For live cell experiments with drug-treatment, 2hr RIF inhibition (100 μg ml^−1^) was performed after the 2hr TetR-EYFP induction, and 1hr SHX (500 μg ml^−1^) treatment was performed during the last hour of TetR-EYFP induction. Live cells after induction or drug treatments were harvested and prepared for imaging as described in the section below. For fixed cell experiments, growth and drug treatments were done as follows: cells were grown to mid-log phase in EZRDM with 0.4% glucose at RT. Cells were treated with drugs when appropriate; SHX treatment was performed at 500 μg ml^−1^ for 1hr, RIF treatment was performed at 100 μg ml^−1^ for 2hr, and novobiocin treatment was performed with 300 μg ml^−1^ for 30 min.

### Sample preparation and imaging conditions

A gel pad made with 3% low-melting-temperature agarose (SeaPlaque, Lonza) in the same growth media (or PBS for fixed cells) was prepared. Live cells were spun-down in a bench-top centrifuge at 8000 rpm for 2 min and resuspended in around 50 μl of fresh growth media (or PBS for fixed cells). An aliquot of 1 μl of the resuspension was then deposited to the agarose gel pad and cells immobilized between the gel pad and a coverslip for imaging as previously described^80,81^. For fixed cell experiments, cells were fixed in 3.7% (v/v) paraformaldehyde (16% Paraformaldehyde, EM Grade, EMS) for 15 min at RT, washed with 1X PBS and imaged immediately. An Olympus IX-81 inverted microscope with a 100X oil objective (UPlanApo, N = 1.4x) was used, with 1.6x additional amplification. Images were captured with an Ixon DU-895 (Andor) EM-CCD with a 13 μm pixel size using MetaMorph (Molecular Devices). Illuminations (405 nm, 488 nm, 561 nm, 647 nm) were provided by solid-state lasers Coherent OBIS-405, Coherent OBIS-488, Coherent Sapphire-561, and Coherent OBIS-647 respectively. Fluorescence was split using a multi dichroic filter (ZT 405/488/561/647rpc, Chroma), and the far-red, red and green channels were further selected using HQ705/55, HQ600/50 and ET525/50 bandpass filters (Chroma). For two-color imaging, the simultaneous, multi-color acquisition was achieved using Optosplit II or Optosplit III (Cairn Research), colored channels were overlaid using calibration images from TetraSpeck beads (Life Technologies, T-7279) as previously described^79,82^, with around 10 nm registration error. Gold fiducial beads (50 nm, Microspheres-Nanospheres, Mahopac, NY) were used to correct for any sample drift during imaging as previously described^83,84^. All superresolution images were acquired with a 10 ms exposure time with ^~^3000-9000 frames. Activation of fluorescent proteins was done simultaneously to fluorophore excitation, and activation laser was kept at a consistent power throughout the imaging session.

### Superresolution imaging data analysis

Molecule localization and fitting of superresolution imaging data were done via thunderSTORM plugin (ImageJ, National Institutes of Health, Bethesda, MD)^85^. Subsequent analysis of localizations was performed using custom Matlab routines. See sections below for data analysis specifics. Custom Matlab routines will be available upon request.

### Blinking correction

To correct for fluorophore blinking and its contribution to false clustering in superresolution images, we utilized a methodology we recently developed, Distance Distribution Correction (DDC)^22^. Briefly, DDC utilizes the finding that the pairwise distance distribution of localizations separated by a frame difference greater than the maximum lifetime of the fluorophore converges upon that of the “true localizations” (not due to the blinking of the emitters). DDC obtains a blinking corrected image by performing a phase search, eliminating localizations of high blinking probabilities, so that the pairwise distance distribution at all frames is consistent with the “true pairwise distance distribution.” We verified that this methodology was significantly more accurate compared to the commonly used thresholding methodology using a variety of simulations and diverse clustering structures, providing the most rigorously scrutinized representation for the locations of the underlying molecules to date.

### Cluster identification

To determine a cluster across the different experimental conditions and different molecular species, we normalized the number of localizations by cell volume so that each cell had the same concentration of localizations. The concentration normalization eliminated the effects of cell size and the noise in detection efficiency from being the dominating factors in the characteristics of the clusters. To do this, we first calculated the volume by outlining the shape of each cell using the outermost localizations to determine an area; this area was then projecting to a 3D volume. We only used cells that had enough localizations (> 700 localizations) to reach the desired concentration threshold for each species.

To obtain the properties of individual clusters, for each species in various conditions, we first eliminated localizations in low-density areas. By calculating the average distance to the closest ten localizations surrounding each localization, we determined whether each localization was in a high-density region if the average distance was greater than a specified value. This calculation was only valid since each cell had the same concentration of molecules, which allowed us to use one defined threshold for each species.

Including only localizations within the high-density regions, we applied a tree-cluster algorithm in MATLAB. Specifically, we utilized the ‘single’ method using the linkage function, which provided us with a tree of hierarchical clusters for the data. We then used the cluster function with a cutoff of 100 nm and the distance criterion. The analysis linked all localizations together as one cluster if they are within 100 nm of each other. As a final step, we only counted clusters that possessed more than a certain percentage of the total localizations. All analysis code will be available upon request.

### Random distribution simulation

To determine whether the clustering of a species was significant, a random distribution of localizations was created, analyzed and compared for each species and condition. To simulate the random distribution of localizations, we first determined the volume of each cell for each condition (as discussed in the cluster determination section). We then randomly placed localizations within this 3D volume according to a uniform distribution; the number of localizations used closely matched to the experimentally collected molecule number for each cell. We then adjusted the concentration of molecules to match the desired concentration used in the cluster determination section and applied the same clustering analyses.

### Colocalization percentage calculation

We calculated the colocalization value from one species’ cluster to the other species’ clusters as the following. For a cluster of species *A* (for example *A*_1_) and the clusters of species B (*B*_1_…*B*_n_), we calculated the pairwise distances between the localizations in (*A*_1_) to the localizations in any *B* (*B*_1_…. *B*_n_) and recorded the shortest distance (*d*_1_). We repeated this calculation for all the other clusters of species *A* (*A*_1_ *…. A*_n_) in the same cell. Therefore, each cluster of *A (A*_1_*…A*_n_) in the cell was associated with a distance (*d*_1_, *d*_2_…*d*_j_). Next, we repeated the same calculation for all clusters of species *A* in all cells and obtained a data structure in which a cluster *i* of species *A* in cell 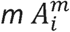 has a distance 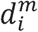. We then selected a threshold distance and counted the number (*n*) of clusters of species *A* that had at least one distance shorter or equal to the threshold distance. Note that we only performed this calculation if both species had clusters within the same cell. The colocalization value of clusters of species *A* to clusters of species *B* was calculated by dividing this number *n* by the total number *(N)* of clusters of species *A* and plotted as a cumulative curve at different threshold distances. As such, a colocalization fraction of 0.8 of RNAP clusters to pre-rRNA clusters at 50 nm means that at 50 nm, 80% of RNAP clusters had at least one pre-rRNA cluster within a distance of 50 nm. Beside the cumulative curves, we also reported colocalization values at a set distance threshold for all experiments conducted in this work (Supplementary Table S3) for ease of comparison between different conditions. Note that the colocalization value from species *A* to species *B* is different from the reverse direction (from species *B* to species *A*) and we reported both in Supplementary Table S3.

### Accounting for experimental cluster detection efficiency

In calculating the colocalization value between two species’ clusters, it is important to consider the detection efficiency of each species’ clusters. Assuming that the detection efficiency for species *B is p*(*p <* 1), and that the true colocalization value from species A to species B is *q*, the measured colocalization value *c* from species A to B will then be modified by the detection efficiency *p* as *c* = *p* · *q.* Therefore, the true colocalization value *q* should be calculated as 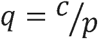.

To determine the detection efficiency of pre-rRNA clusters for the rich medium growth condition, we used two L1 probes with the same sequence but different dye labels (Alexa Fluor 488 and 647, respectively) to hybridize with pre-rRNAs in the same cells. Because the probe sequences were the same, pre-rRNA clusters identified by the two colors should be identical and colocalize with each other 100%. Therefore, the lower than 100% colocalization value we detected from one color to the other, likely resulting from dye properties and cluster thresholding, allowed us to calculate the detection efficiency of the L1 probe. As shown in Supplementary Fig. S6a, we observed nearly identical cumulative curves (After blinking correction) of the colocalization value from L1-Alexa Fluor 488 to L1-Alexa Fluor 647 and *vice versa.* At 50 nm, the detection efficiency of both probes was approximately 80%.

To determine the detection efficiency of RNAP clusters, we used a computational approach (Supplementary Fig. S7) due to the inability to obtain a fully functional RNAP-Dronpa-PAmCherry tandem dimer fusion on the chromosome. In the computational approach, we randomly split into two channels RNAP localizations of cells that had at least twice the predefined concentration of localizations, so that there were two sets of localizations in a cell with the desired concentration, mimicking cluster detection using two different colors. We then performed the cluster analysis on each set of localizations and calculated the colocalization value between the clusters identified in the two sets at different threshold distances (Supplementary Fig. S7). We further verified this computational approach using the experimentally measured colocalization curves of the L1 probes of two different dyes and obtained the same result (Supplementary Fig. S6a).

The colocalization cumulative curves between the two sets of clusters provided us with the best possible colocalization at each distance given our detection efficiency. In all colocalization curves reported in this work except for Supplementary Fig. S6a and S7, we adjusted the colocalization values by dividing the measured colocalization value by the measured detection efficiency value at the same distance.

### smFISH - L1 probe labeling of pre-rRNA

We performed smFISH using a previously published protocol^80,86^. Briefly, cells were grown in EZRDM glucose as previously described; 5 ml of mid-log phase cells were fixed with 3.7% (v/v) paraformaldehyde (16% Paraformaldehyde, EM Grade, EMS), placed for 30 min on ice. Next, cells were harvested via centrifugation, and subsequently washed two times in 1X PBS. Cells were then permeabilized by resuspending in a mixture of 300 μl of H_2_O and 700 μl of 100% ethanol and incubating with rotation at RT for 30 min. Cells were stored at 4 °C until next day. Wash buffer was freshly prepared with 40% formamide and 2x SSC and put on ice. Cells were spun-down in a bench-top centrifuge at 10000 rpm for 3 min and the cell pellet was resuspended in 1 ml of wash buffer. The sample was placed on a nutator to mix for 5 min at RT. Hybridization solution was prepared with 40% formamide and 2x SSC, subsequently, dye-labeled oligo probes were added to hybridization solution to a final concentration of 1 μM. Cell were spun-down again and 50 μl of hybridization solution with probe was added to the pellet. The hybridization sample was mixed well and placed overnight in a 30 °C incubator. Next day, 10 μl of hybridization sample was washed with 200 μl of fresh wash buffer and incubated at 30 °C for 30 min, this was repeated one more time. The washed sample was imaged immediately: without STORM imaging buffer for ensemble fluorescence, with STORM buffer to induce dye blinking for superresolution imaging, glucose oxidase + thiol STORM buffer was used to image samples with only dye labeling (50 mM Tris (pH 8.0), 10 mM NaCl, 0.5 mg ml^−1^ glucose oxidase (Sigma-Aldrich), 40 μg ml^−1^ catalase (Roche), 10% (w/v) glucose and 10 mM MEA (Fluka))^87^. Thiol only STORM buffer (10 mM MEA, 50 mM Tris (pH 8.0), 10 mM NaCl) was used to image samples with both endogenously expressed fluorescent proteins and dye labeling. This was to preserve the fluorescent signal from fluorescent proteins, since the presence of glucose oxidase in the STORM buffer tended to quench the fluorescent protein signal.

Pre-rRNA transcripts were detected with a single probe L1, conjugated at the 5’ with either Alexa Fluor 488 (NHS ester) or Alexa Fluor 647 (NHS ester) (IDT) (Supplementary Fig. S5)^25^. Upon receiving the commercial oligos, a working stock (50 μM) was made and aliquoted for storage at −20 °C.

### Image processing of smFISH ensemble fluoresence images

Ensemble intensity measurements were performed using ImageJ (National Institutes of Health, Bethesda, MD). Ensemble fluorescence images with focus plane at mid-cell were segmented manually using their corresponding bright-field images. Each cell’s total fluorescent intensity was calculated as: (area of the segmented cell * (mean intensity inside cell – mean intensity of background region)). Around 50-100 cells were used to represent the total cellular fluoresence intensities for a single experimental condition. See corresponding figure captions for the exact number of cells used in the calculations.

### DNA staining in fixed cells using Hoechst dye (33342)

Hoechst dye (bisBenzimide H33342 trihydrochloride) was used to stain chromosomal DNA in *E. coli* cells. Stained cells were subsequently visualized via 3D SIM on a GE OMX SR SIM scope, with a 60x objective. Briefly, cells were grown to mid-log phase (O.D._600_ ^~^ 0.3-0.4) in EZRDM at RT. For all conditions except for Δ6*rrn*, the strain RpoC-PAmCherry was used for DNA staining and considered as the wt strain. Hoechst dye (0.5 μl of 10 mg ml^−1^ stock) was added during the last 10 min of cell growth. After 10 min of incubation with Hoechst dye, 1 ml of the liquid culture was immediately harvested and spun-down in a bench-top centrifuge at 8000 rpm for 2 min and the cell pellet was washed with 1 ml of 1x PBS. For fixation, the cell pellet was resuspended in 1 ml of 3% paraformaldehyde in 1x PBS, placed on a nutator and fixed at RT for 15 min. After fixation, cells were subsequently spun-down at 8000 rpm for 2 min and the cell pellet was washed with 1 ml of 1x PBS. About 35 μl of 1x PBS was used to resuspend the cell pellet as a final step before mounting. An equal volume of fixed cells in PBS and anti-fading buffer (60% glycerol, 20% NPG (n-propyl gallate, 1x PBS) was combined to a total of 50 μl and used for mounting between Poly-L-Lysine treated coverslip and cover glass. Excess liquid was siphoned away with a Kimwipe tissue, and the coverslip was sealed on the glass slide using clear nail polish. Imaging was performed 30-60 min after sample preparation.

3D SIM Imaging conditions were as follows: 5% laser power for 405 nm laser excitation, 30 ms exposure time, with standard 125-nm interval Z-sections, and a pixel size of 40 nm. Images were collected using the standard SRx software and reconstructed using standard SIM reconstruction parameters.

For a more quantitative comparison between different experimental conditions, we calculated the nucleoid territory occupancy in cells. Briefly, we used intensity thresholding to isolate both the cell area voxels (a lower intensity threshold), and the DNA area voxels (a higher intensity threshold) and used (DNA area/cell area) to calculated the percentage of total cell area that was occupied by DNA, a representative 15 cells were used for this calculation for each experimental condition. Additionally, we constructed 2D histograms of the DNA signal intensities for each condition. The DNA intensity from the projected Z-stack of the eight 125-nm Z slices for each cell were combined for each condition, 15 cells were used for each experimental condition, the cells were rotated, and the long axis was normalized to each cell’s cell length; the 2D histograms were represented in a standard 1 μm × 3 μm cell.

